# Transmembrane Stem Factor Nanodiscs Enhanced Revascularization in a Hind Limb Ischemia Model in Diabetic, Hyperlipidemic Rabbits

**DOI:** 10.1101/2023.03.20.533550

**Authors:** Eri Takematsu, Miles Massidda, Gretchen Howe, Julia Goldman, Patricia Felli, Lei Mei, Gregory Callahan, Andrew D Sligar, Richard Smalling, Aaron B. Baker

**Author notes:** Correspondence to: Aaron B. Baker, Ph.D. University of Texas at Austin, Department of Biomedical Engineering, 1 University Station, BME 5.202D, C0800, Austin, TX 78712.

## Abstract

Therapies to revascularize ischemic tissue have long been a goal for the treatment of vascular disease and other disorders. Therapies using stem cell factor (SCF), also known as a c-Kit ligand, had great promise for treating ischemia for myocardial infarct and stroke, however clinical development for SCF was stopped due to toxic side effects including mast cell activation in patients. We recently developed a novel therapy using a transmembrane form of SCF (tmSCF) delivered in lipid nanodiscs. In previous studies, we demonstrated tmSCF nanodiscs were able to induce revascularization of ischemia limbs in mice and did not activate mast cells. To advance this therapeutic towards clinical application, we tested this therapy in an advanced model of hindlimb ischemia in rabbits with hyperlipidemia and diabetes. This model has therapeutic resistance to angiogenic therapies and maintains long term deficits in recovery from ischemic injury. We treated rabbits with local treatment with tmSCF nanodiscs or control solution delivered locally from an alginate gel delivered into the ischemic limb of the rabbits. After eight weeks, we found significantly higher vascularity in the tmSCF nanodisc-treated group in comparison to alginate treated control as quantified through angiography. Histological analysis also showed a significantly higher number of small and large blood vessels in the ischemic muscles of the tmSCF nanodisc treated group. Importantly, we did not observe inflammation or mast cell activation in the rabbits. Overall, this study supports the therapeutic potential of tmSCF nanodiscs for treating peripheral ischemia.

## Introduction

Diabetes mellitus affects approximately 350 million people, leading to the death of an estimated 4.6 million people in the world per year^1,2^. Diabetes leads to a disturbance of the blood vessel by promoting vascular inflammation and endothelial cell dysfunction^3,4^. These abnormalities increase the severity of vascular disease in diabetic patients^5^. As a complication of diabetes, 30 to 40 percent of patients age 50 and older develop peripheral artery disease (PAD)^6^. Severe PAD increases the risk of non-healing ulcers, pain from intermittent claudication, and worst case for limb amputation. Current standard cares for PAD include physical therapy, medication, and surgical revascularization. Surgical bypass is an important treatment option for sever PAD, however many patients especially elderly patients are not eligible because of their diffusive arterial occlusions, no suitable veins for grafting and comorbidity^7^. Therapeutic angiogenesis by growth factors or growth factor genes is an appealing strategy to treat ischemia in this context but has been difficult to translate in the clinical setting. Many growth factors/gene therapies have shown promise in preclinical studies for ischemia only to show disappointing results in clinical trials in human patients.^8–10^

Stem cell factor (SCF) acts as a hematopoietic cytokine that communicates via the c-Kit receptor (CD117) and is also recognized as Kit ligand, Steel factor, or mast cell growth factor^11^. SCF is produced in cells as a transmembrane protein through alternative splicing, which is then enzymatically divided into soluble SCF or a shorter version without the cleavable domain, remaining membrane-bound as transmembrane SCF (tmSCF)^12^. The activation of c-Kit through SCF signaling is crucial for maintaining hematopoietic stem cells (HSCs) and progenitor cells in the bone marrow^13,14^. Numerous potential therapeutic applications for SCF exist, such as enhancing the survival and expansion of HSCs after radiation exposure^15,16^, promoting neuroprotection following a stroke^17–19^, and aiding the heart’s recovery after a myocardial infarction^20^. In addition, SCF plays a crucial part in controlling mast cell development and activation^21,22^. Administering SCF results in mast cell activation and anaphylaxis in both animal studies and clinical trials, significantly restricts its potential for therapeutic use^22–26^. We recently found that tmSCF delivered in proteoliposomes or lipid nanodiscs was effective in inducing revascularization of ischemic limbs in diabetic and wild type mice but did not lead to mast cell activation^27^. Thus, tmSCF may provide benefits of soluble SCF without the toxic side effects that limited its usefulness as a therapeutic.

While our previous studies have supported the potential of tmSCF nanodiscs as therapies for ischemia, large animal studies are needed to demonstrate efficacy to support further studies in human trials. For peripheral ischemia, rabbits are typically used as the precursor to human studies for treating PAD. Our group recently developed an optimized rabbit model of hindlimb ischemia that includes hyperlipidemia and diabetes^28,29^. A major advantage of this model is that it exhibits resistance to angiogenic therapies and longer ischemia, similar to human patients, in contrast to health rabbit models that revascularize more quickly. In this study, we used this advanced rabbit model to evaluate the efficacy of tmSCF nanodiscs in treating peripheral ischemia. Overall, our results demonstrate that tmSCF nanodiscs improve revascularization in ischemia compared to alginate gel control without significant inflammatory reactions.

## Methods

### Preparation of tmSCF Nanodiscs

1-palmitoyl-2-oleoyl-sn-glycero-3-phosphocholine (POPC) was stored in chloroform, so we first removed chloroform by rotary evaporator. After that, POPC was resuspended in sodium cholate (100 mM). MSP protein was then added to phospholipid solution, and the detergent concentration was adjusted between 14 and 40 mM. This construct was incubated for 15 min at 4 °C. To solubilize the membrane protein, tmSCF was incubated in the n-octyl-β-D-glucopyranoside (1% w/v) for 15 min at 4 °C. After 15 min incubation of lipid construct and tmSCF protein, these were combined and incubated for 1 hour at 4 °C. Final detergent concentration was adjusted to 20 mM with sodium cholate. Finally, detergent was removed by dialysis and biobeads (**Supplemental Fig.1**).

### Preparation of Alginate Gels and Crosslinking

Sterile alginate powder (Sigma) was added to sterile saline to create a 2% w/v solution. The final concentration of tmSCF nanodiscs was adjusted to 50 μg/ml in this 2% alginate solution. To prepare the crosslinker, calcium sulfate was added to sterile saline to create 0.2% w/v solution. Both the alginate solution and the crosslinker were taken up into a 1 ml syringe just before injection (100 μl of each solution for 200 μl total per injection).

### Induction of Diabetes in Rabbits

Studies involving animals were performed with the approval of the University of Texas at Austin and the University of Texas Health Science Center at Houston Institutional Animal Care and Use Committees (IACUCs), the Animal Care and Use Review Office (ACURO) of the United States Army Medical Research and Materiel Command Office of Research Protections, and in accordance with NIH guidelines for animal care. New Zealand rabbits were transitioned from standard alfalfa chow to a 0.1% cholesterol diet over the course of five days. After two weeks on the 0.1% cholesterol diet, rabbits were induced with diabetes using an intravenous alloxan injection. After sedating the rabbits, a bassline blood glucose measurement was attained. Alloxan (100 mg/kg) was injected through an IV into the rabbit at flow rate of 1 ml/min for eight minutes using a syringe pump. Blood glucose levels were monitored closely for 12 hours after the injection. A successful induction of diabetes was determined if the rabbit’s blood glucose level remained over 150 mg/dl prior to insulin administration.

### Hind limb Ischemia Surgery and Treatments

To induce aischemia in the hind limb of New Zealand rabbits, a longitudinal incision was made in the skin over the femoral artery. The femoral artery was exposed using blunt dissection. One percent lidocaine was applied to the area to reduce nerve irritation and promote vasodilation. Continued blunt dissection was used to expose the entire length of the femoral artery and branches including the inferior epigastric, deep femoral, lateral circumflex, and superficial epigastric arteries. The tissue was kept moistened with saline to avoid damage. The femoral artery was then carefully separated from vein and nerve. The femoral artery was then ligated with 4.0 silk sutures, cut, and excised. Two weeks after hind limb ischemia surgery, ten syringes containing 100 μl of alginate with treatment and 100 μl of calcium sulfate crosslinker were prepared for intramuscular injection (200 μl total volume of injection). Treatments included tmSCF nanodiscs in alginate and an alginate-only control.

### Angiography Quantification

Angiograms were quantified using grid analysis techniques with ImageJ Software. Briefly, brightness and contrast were adjusted to better visualize vessels. A grid overlay of 100 pixels per square was used for foot quantification and 750 pixels per square was used for the thigh vasculature quantification. The multi-point tool was used to count intersections between the grid and vessels. To access the carotid artery, an incision was made just lateral to the trachea. Blunt dissection was used to expose the carotid artery and separate it from the jugular vein and vagus nerve. Ligatures were placed at the proximal and distal ends of the carotid artery. The distal end was tied off and a ligaloop was placed at the proximal end. A wire insertion tool was then inserted into the artery. Using the tool, a guidewire was fed into the artery to the aortic bifurcation in the descending aorta. The insertion tool was then removed and a 3F pigtail angiographic catheter was placed over the wire and advanced 2 cm proximal to the aortic bifurcation. Nitroglycerine and lidocaine were given to increase vasodilation. Contrast media was injected through the catheter using an automated angiographic injector. Angiography was performed before femoral ligation, after femoral ligation, and before sacrifice at week 10.

### Statistical Analysis

All results are shown as mean ± standard error of the mean. Comparisons between only two groups were performed using a 2-tailed Student’s t-test. Differences were considered significant at p<0.05. Multiple comparisons between groups were analyzed by 2-way ANOVA followed by a Tukey post-hoc test. A 2-tailed probability value <0.05 was considered statistically significant.

## Results

### Transmembrane SCF nanodiscs enhance revascularization of the ischemic thigh muscles of diabetic, hyperlipidemic rabbits

As optimized in our previous studies, rabbits were given a high fat diet for four weeks and were induced with diabetes for two weeks using alloxan prior the surgery to induce limb ischemia^28,29^. The femoral artery was ligated in the right leg of the rabbits and an angiogram was taken before and immediately after the surgery to assess the induction of ischemia in the limbs (**Supplemental Fig. 2**). The rabbits were then allowed to recover for two weeks to avoid treatment during the acute healing phase of recovery^29^. The rabbits were given ten injections of alginate with or without the tmSCF nanodiscs and recovery was assess using angiography after 10 weeks (**Fig. 1A; Supplemental Fig. 2**). These angiograms were analysed quantitatively for several measures of vascularity. A grid was superimposed onto the angiograms and intersections between the grid and vessels were counted. Vessel grid intersection counts showed significant vasculature increase for the tmSCF nanodisc group compared to the alginate control (**Fig. 1B**). When ratioed to the contralateral control limb at week 10, the tmSCF nanodisc group significantly improved recovery compared to the alginate control (**Fig. 1C**). A ratio was also taken of the grid intersections counted in the ischemic thigh at week 10 to the intersections counted in the ischemic thigh before hind limb surgery. The results show a significant increase in vascularity for the tmSCF nanodisc group as well (**Fig. 1D**). A new vessel formation is indicated by tortuosity (corkscrew like morphology), which was also counted to assess angiogenic capacity of tmSCF nanodiscs. The results also showed a significant increase in tmSCF nanodisc group in comparison to control. (**Fig. 1E**).

**Figure 1.**
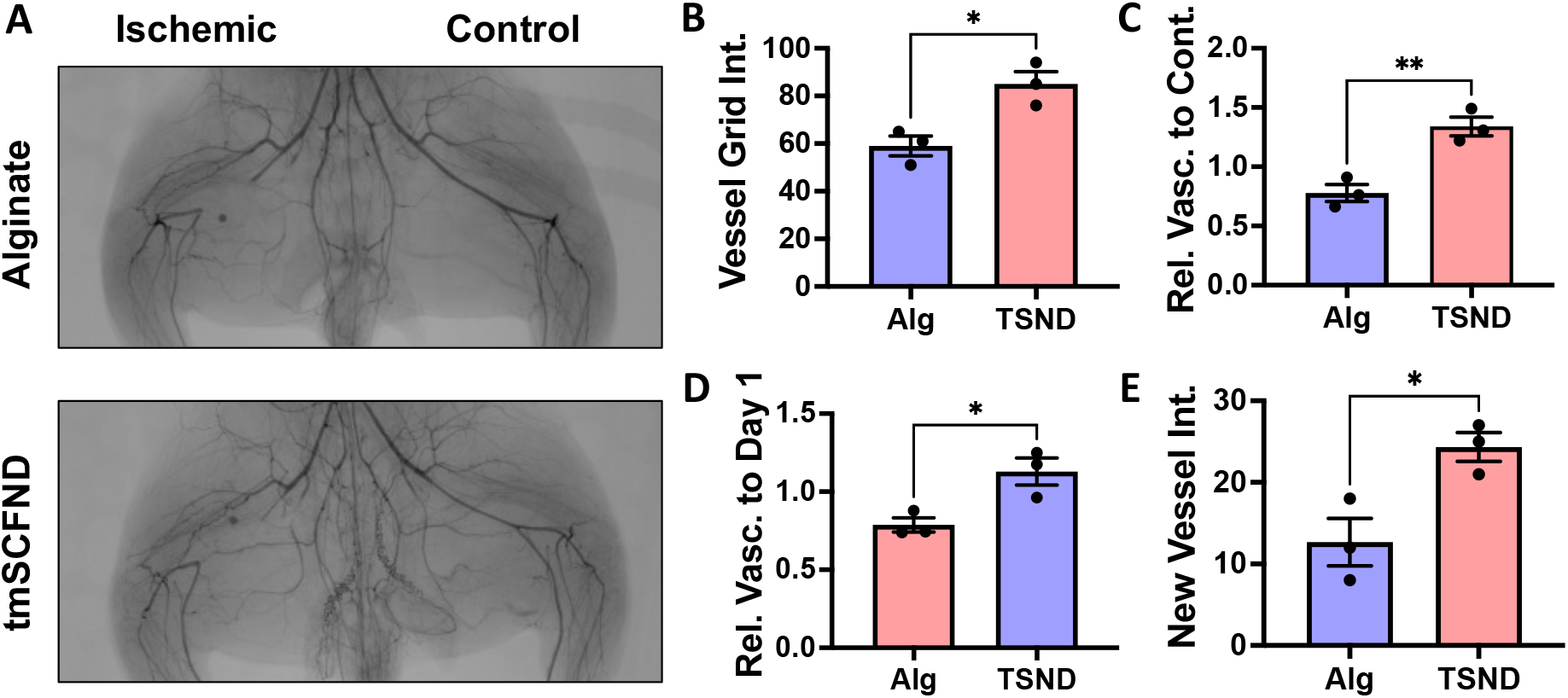
Transmembrane SCF nanodiscs enhance revascularization in the thigh of diabetic, hyperlipidemic rabbits with hindlimb ischemia. (A) Angiograms of the thighs of the rabbits at week 10. The control limb is shown on the right and the limb with the ligated femoral artery is on the left. (B) Quantification of vessel grid intersections counted in the ischemic thigh at the model endpoint. The vessels are counted as number of intersections with an overlayed grid. (C) Relative vascularity of the ischemic thigh ratioed to the contralateral control thigh at the model endpoint. (D) Relative vascularity of the ischemic thigh ratioed to the thigh at day 1 prior to ligation. (E) Quantification of new vessels counted in the ischemic thigh at the model endpoint. \**p* < 0.05 vs. alginate; \*\**p* < 0.01 vs. alginate. (n=3).

### Transmembrane SCF nanodiscs enhance revascularization of the ischemic calf and foot of diabetic, hyperlipidemic rabbits

Postoperative angiogram showed the most severe ischemia is the area of the foot and calf muscles (**Supplemental Fig. 2**). Ten weeks after the operation, robust vascular network formation were observed at the foot of the tmSCF nanodiscs treated group while almost few or no blood vessels was observed in the control group by angiography (**Fig. 2A**). Quantitative analysis on vessel count showed a significantly higher blood vessel numbers in tmSCF nanodisc group in comparison to the control (**Fig. 2B**). Similarly, significantly higher ratio of vascular count at the ischemic calf and foot when ratioed to the contralateral control at week 10 (**Fig. 2C**). A ratio of vessel counts at the ischemic thigh at week 10 to the vessel counts before surgery also demonstrated the effectiveness of the tmSCF nanodisc treatment (**Fig. 2D**).

**Figure 2.**
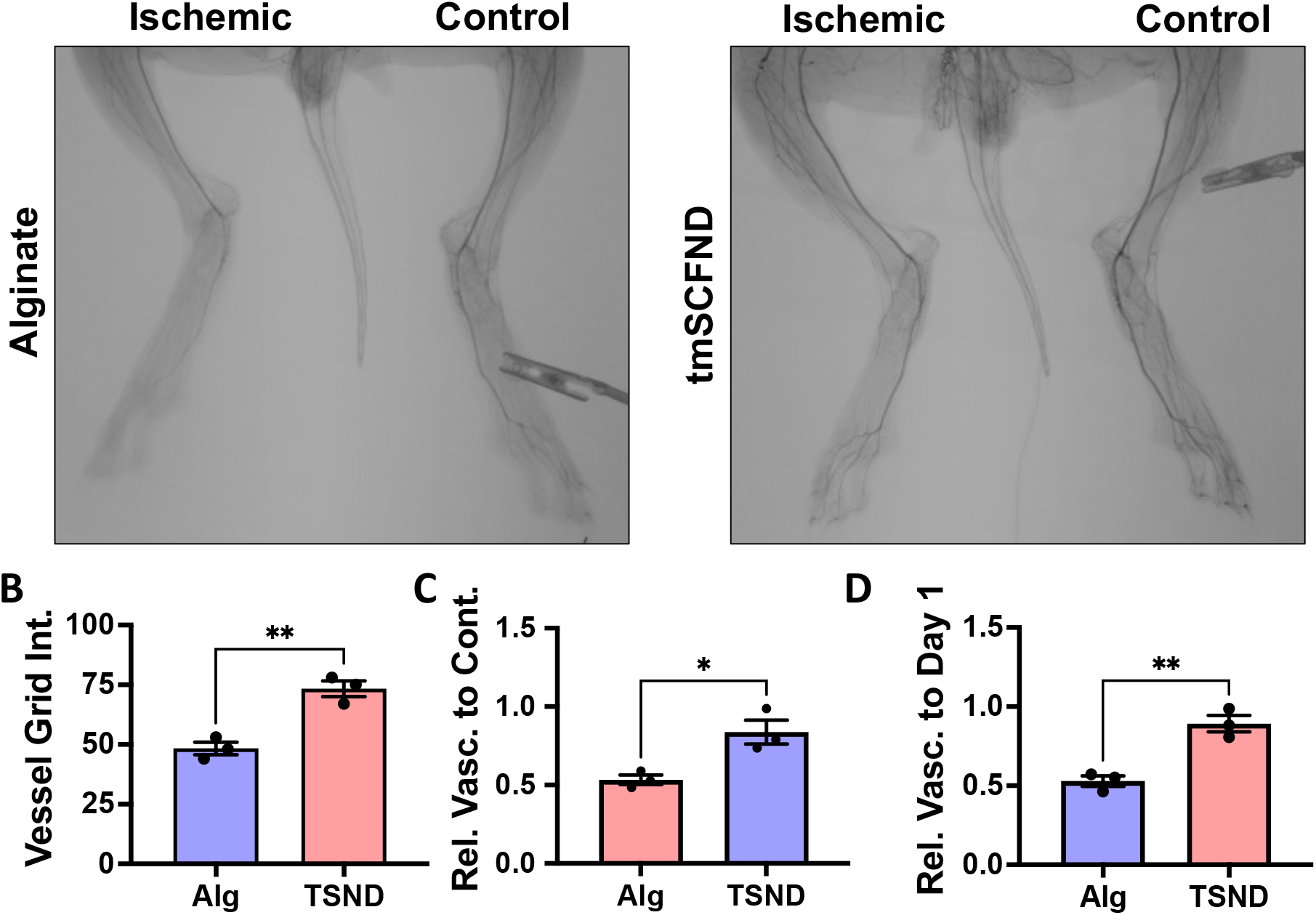
tmSCFND enhances revascularization in the calf and foot of diabetic, hyperlipidemic rabbits with hindlimb ischemia. (A) Angiograms of the lower limb of the rabbits at week 10. (B) Quantification of vessels counted in the ischemic calf and foot at the model endpoint. The vessels are counted as number of intersections with an overlayed grid. (C) Relative vascularity of the ischemic limb ratioed to the contralateral control limb at the model endpoint. (D) Relative vascularity of the ischemic calf and foot ratioed to the calf and foot at day 1 prior to ligation. \**p* < 0.05 versus alginate; \*\**p* < 0.05 versus alginate. (n=3)

### Histological analysis confirmed increased vascularity in tmSCF nanodisc treated leg of rabbits with diabetes and hyperlipidemia

We next performed a histological analysis on biopsies from the ischemic muscle of the rabbits. PECAM immunostaining was used to count small and large vessels formation (**Fig. 3A**). Significantly more small vessels and large vessels were confirmed with tmSCF nanodisc treatment, indicating that tmSCF nanodiscs induced both angiogenesis and arteriogenesis in the rabbit ischemia model (**Fig. 3B**). Importantly, no signs of inflammation was found on the H&E staining of thigh and calf muscle tissues.

**Figure 3.**
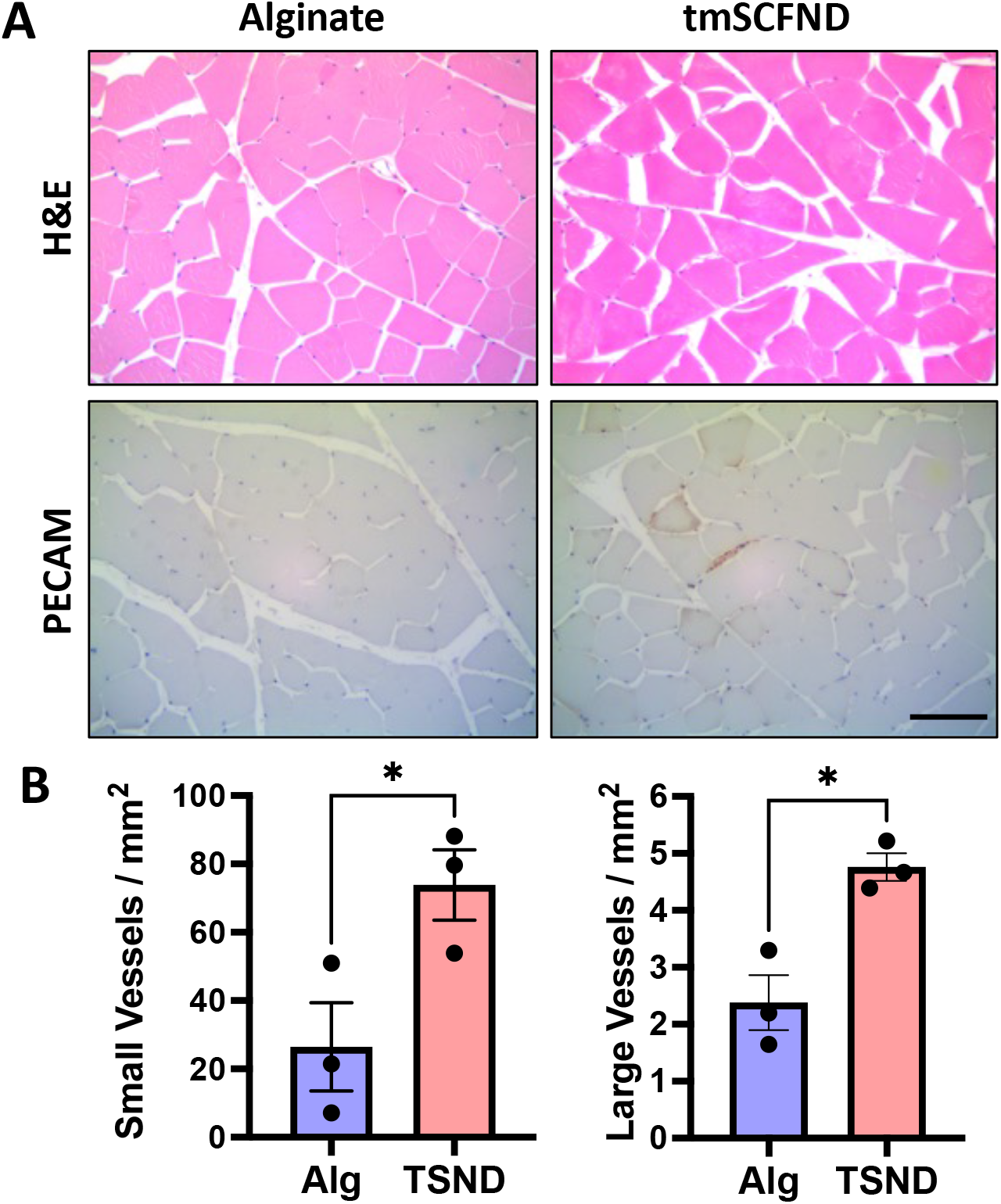
Histological analysis of vascularization of calf treated by tmSCF nanodiscs. (A) Upper: H&E staining analysis of biopsies from the ischemic hind limbs of rabbits. Bottom: PECAM staining analysis of biopsies from the ischemic hind limbs of rabbits. Scale bar = 100 μm. (B) Left: Quantification of small vessels counted in the ischemic calf at week 10. Right: Quantification of arterioles vessels counted in the ischemic calf at week 10. *p < 0.05 versus alginate group (n = 3).

## Discussion

Stem cell factor is a therapeutic protein with many potential uses for treating disease. Unfortunately, its use has been severely limited by toxic effects that were observed in animal studies and human clinical trials^22–26^. Managing ischemia in diabetic individuals has presented continuing challenges in the clinical setting, particularly for those undergoing bypass operations or percutaneous procedures^30,31^ where diabetic patients have worse outcomes and reduced benefits from therapy and interventions. Our previous work demonstrated that tmSCF nanodiscs enhance revascularization in healthy and diabetic mice^32^. In this work, we evaluated the efficacy of tmSCF embedded in nanodiscs for therapeutic angiogenesis using an optimized rabbit model of hindlimb ischemia with diabetes and hyperlipidemia^28,29^. Angiography of the thigh and calf muscle treated with tmSCF nanodiscs showed significantly higher numbers of functional blood vessels and new vessels. Histological analysis on PECAM staining and H&E staining also demonstrated significantly higher numbers of small and large vessel formations compared control, indicating that tmSCF nanodiscs support both angiogenesis and arteriogenesis. The findings are particularly significant given the rabbit model’s resistance to angiogenesis and longer-term ischemia^28,29^. Finally, we did not observe any signs of mast cell activation on histological analysis, consistent with our prior studies in mice^27^.

Prior studies have explored the use of many treatments in rabbits and other large animals for treating peripheral ischemia and ischemia wounds^33,34^. A lack of uniformity in the techniques employed for performing and analyzing preclinical hindlimb ischemia models in rabbits makes it difficult to directly compare with other studies. In addition, the majority of prior studies have also used healthy rabbits, which have more rapid recovery from ischemia and responsiveness to angiogenic treatments. Overall, the improvement in revascularization observed in our research using a diseased rabbit model is comparable or superior to earlier studies that utilized healthy rabbit models of hindlimb ischemia to assess protein therapeutics^35–57^, cell therapies^58–62^, or gene therapy^55,63–75^. Consequently, our work supports that tmSCF nanodiscs would have potential as a therapeutic for ischemia, even in the context of therapeutic resistance as would be found in many human patients that have hyperlipidemia and diabetes.

## Declarations

### Ethics approval and consent to participate

Studies involving animals were performed with the approval of the University of Texas at Austin and the UTHealth Science Center at Houston Institutional Animal Care and Use Committee (IACUC), the Animal Care and Use Review Office (ACURO) of The United States Army Medical Research and Materiel Command Office of Research Protections, and in accordance with NIH guidelines for animal care.

### Consent for publication

Not applicable.

### Availability of data and materials

The datasets used and/or analyzed during the current study are available from the corresponding author on reasonable request.

### Authors’ contributions

E.T. analyzed data and wrote/edited the manuscript. M.M., G.H., J.G., P.F., L.M., G.C., A.D.S. and R.S. performed experiments, data collection and analysis. A.B.B. conceived the original idea, analyzed experiments, and wrote/edited the manuscript.

## Acknowledgements

The authors gratefully acknowledge funding through the DOD CDMRP (W81XWH-16-1-0580; W81XWH-16-1-0582) and the National Institutes of Health (1R01HL141761-01) to ABB.

## Supplemental Figures

**Supplemental Figure 1.**
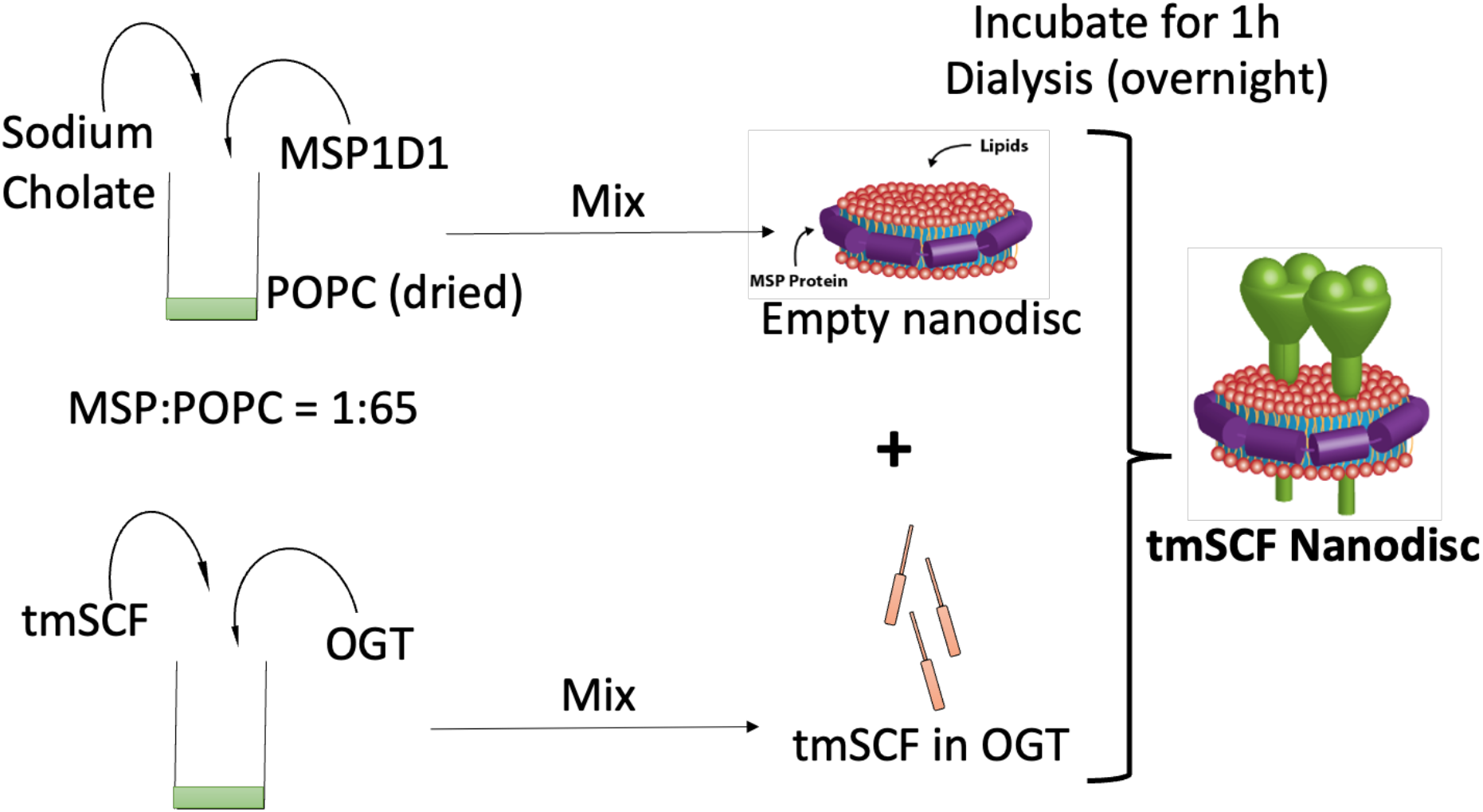
Schematic illustration for fabricating tmSCF nanodiscs.

**Supplemental Figure 2.**
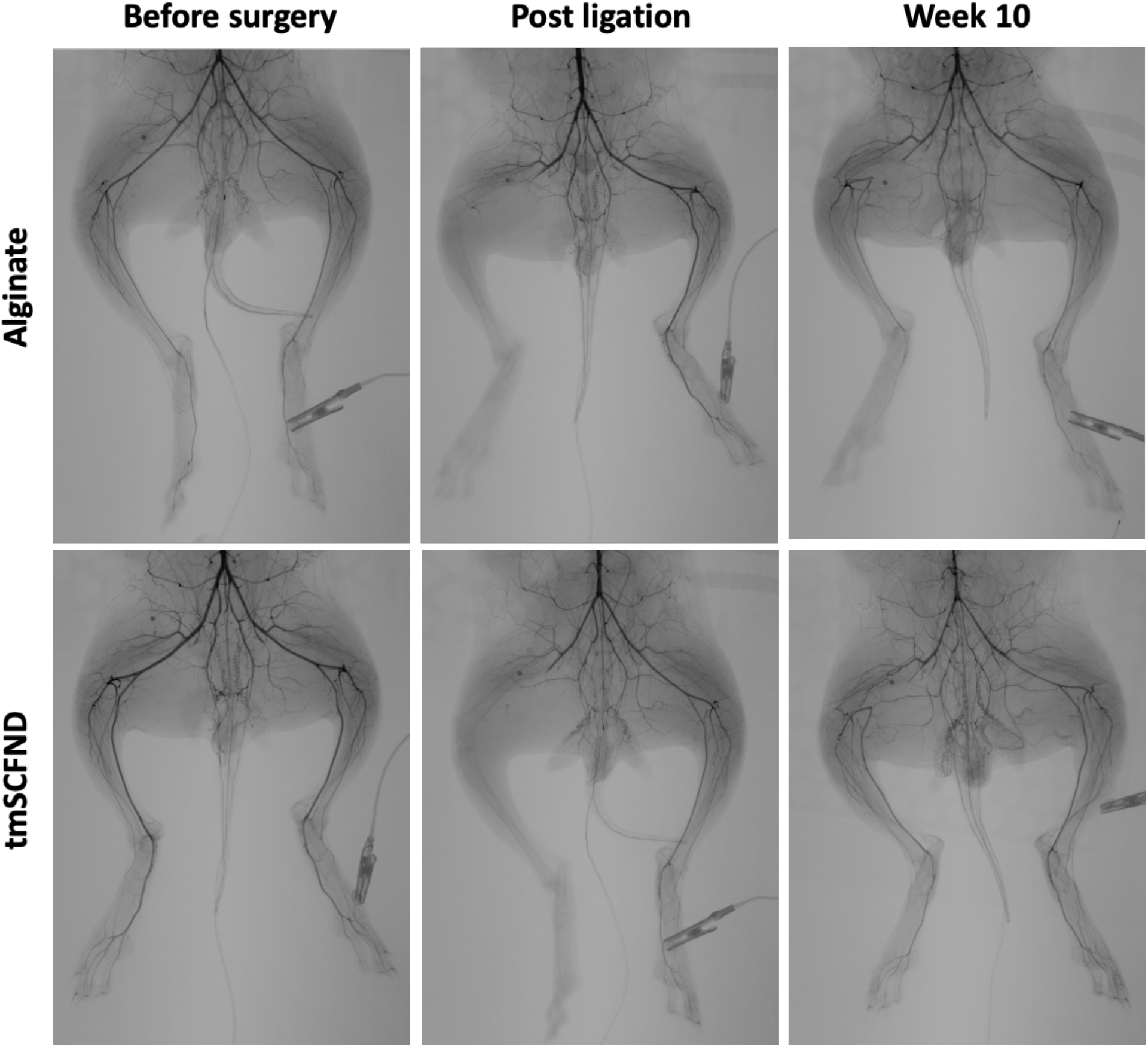
Full angiograms for the rabbits before/after ligation and 10 weeks postsurgery.

